# Cerebrovascular Responses to Static and Rhythmic Handgrip Exercises

**DOI:** 10.1101/2025.09.11.675716

**Authors:** Elric Y. Allison, Yixue Mei, Huseyn A. Ismayilov, Geoff B. Coombs, Jeremy J. Walsh, Michael J. Carter, Trevor King, Baraa K. Al-Khazraji

## Abstract

Handgrip exercise (HGE) is a safe, accessible exercise modality shown to improve cardiovascular health and is particularly promising for individuals with limited mobility who cannot engage in traditional exercise. Given that contraction type and intensity influence systemic cardiorespiratory variables that affect cerebral blood flow regulation, this study examined the acute systemic hemodynamic and cerebrovascular responses to static and rhythmic HGE protocols at varying intensities. Thirty-three healthy young adults (17 males; 16 females age 22(1) years) performed four separate 5-minute HGE protocols in a randomized order: static HGE at 15% (S15) of maximal voluntary contraction (MVC), static HGE at 30% MVC, rhythmic HGE at 30% MVC, rhythmic HGE at 60% (R60) MVC. We hypothesized that rhythmic HGE at higher intensities would produce the greatest cerebrovascular responses due to enhanced venous return and cardiac output, while static HGE at higher intensities would elicit the greatest systemic (i.e., blood pressure, HR, ventilation) responses. Cerebral (middle cerebral artery blood velocity [MCAv] and cerebrovascular conductance index [MCA_CVCi_], internal carotid artery [ICA] diameter, velocity, blood flow, and shear rate) and systemic hemodynamics (systolic [SBP], diastolic [DBP], mean arterial pressure [MAP], heart rate [HR], cardiac output [CO]), and end-tidal carbon dioxide (P_ET_CO_2_) levels were averaged over the final 30s of each minute of exercise. There was a significant time and protocol interaction effect on HR (*p*<0.001). We found significant main effects of exercise protocol for MCAv (*p*<0.001), MCA_CVCi_ (*p*<0.001), ICA diameter (*p*=0.01) and blood flow (*p*=0.001), and P_ET_CO_2_ (*p*=0.002). Greatest increases in MCAv alongside the largest reduction in ICA blood flow occurred in R60. The greatest increase in MCA_CVCi_ and ICA blood flow (from baseline) occurred in R30 compared to other protocols. In addition to the greater cerebrovascular responses, we also observed more modest systemic responses (lower HR, MAP, CO) and lower self-reported ratings of perceived exertion (*p*<0.001) in R30 compared to other protocols. Acute increases in MCA_CVCi_ and ICA blood flow observed in R30 (despite the lower perceived effort) may suggest that rhythmic HGE at low-moderate intensities can be a tolerable prescription for inducing exercise-related cerebrovascular adaptations (i.e., improved cerebral perfusion) in populations that may require adapted physical activity.

## Introduction

Cerebral blood flow (CBF) is essential for sustaining neuronal metabolism and long-term brain health (1). Repeated exposure to elevations in vascular shear stress is thought to elicit adaptive responses that underpin the cerebrovascular adaptations to exercise and support the brain health benefits of regular physical activity (2). Absence of exercise-related increases in shear stress on the cerebral arteries may lead to cerebrovascular dysfunction (3) and hypoperfusion (4) with implications on long term brain health.

Despite the clear metabolic and functional benefits of traditional aerobic and resistance exercise, alternative exercise modalities are necessary to promote physical activity among individuals with reduced functional capacity or limited mobility. Handgrip exercise (HGE) has been endorsed by the American Heart Association as an effective intervention for blood pressure management (5). Cross-sectionally, handgrip strength is a strong predictor of metabolic health (6) cardiovascular health (7), long-term brain health (8,9), and all-cause mortality (10). The practicality, low barrier to entry, safety, and minimal equipment requirements makes HGE an attractive alternative option to more traditional and physically demanding forms of exercise.

Findings from studies using HGE and other exercises (ie., quadriceps extension) suggests that both exercise intensity and muscle contraction type (i.e., static vs rhythmic) influence systemic parameters critical to CBF regulation, including sympathetic nerve activity (11), blood pressure (12), and total peripheral resistance and cardiac output (CO) (13). Variations in HGE protocols make it challenging to determine an optimal and tolerable HGE protocol for targeting the cerebral vasculature to promote adaptation. Static HGE has been suggested to produce greater increases muscle sympathetic nerve activity (MSNA) compared to rhythmic HGE (11), in part due to metabolite accumulation and subsequent augmented stimulation of the metaboreflex (14). During static HGE, stimulation of the metaboreflex elicits sympathetic activation via group III/IV afferents and subsequently increases total peripheral resistance and blood pressure in an intensity-dependent manner (15,16). Compared to rhythmic HGE, greater blood pressure responses during static HGE may elicit compensatory autoregulatory mechanisms, attenuating the effects of blood pressure on the CBF response to exercise (1). In contrast, the lower sympathetic and pressor response to rhythmic HGE (11,12) may be related to intermittent relaxation phases that allow for more effective metabolite clearance (17). Additionally, rhythmic contractions facilitate venous return via the muscle pump (18), supporting increased cardiac output which may further elevate CBF during exercise (19). While the underlying mechanisms require further elucidation, the discrepant physiological profiles of static and rhythmic HGE may have important implications for cerebrovascular regulation and long-term brain health through adapted physical activity.

Despite the accessibility and potential benefits of HGE, no study to date has systematically compared the cerebrovascular and systemic hemodynamic responses across commonly used HGE protocols. In particular, the influence of different HGE protocols on CBF and systemic physiological responses remains unclear. Understanding these relationships is essential to optimize small muscle mass exercise and may be particularly useful for individuals who face physical limitations and cannot engage in more dynamic forms of exercise. The primary purpose of this study was to examine the acute cerebrovascular (middle cerebral artery blood velocity [MCAv] and conductance index [MCA_CVCi_], internal carotid artery [ICA] diameter, velocity, blood flow, and shear rate) responses to different HGE protocols in healthy young adults, considering both contraction type and exercise intensity. We also investigated the effects of different HGE protocols on systemic physiological factors that influence cerebral hemodynamics, including heart rate (HR), blood pressure, cardiac output (CO), and end-tidal carbon dioxide (P_ET_CO_2_). We hypothesized that higher intensity rhythmic HGE would produce the greatest increase in MCAv, MCA_CVCi_, and ICA blood flow, while higher intensity static HGE would elicit the greatest systemic (ie., blood pressure, HR, ventilation), responses compared to each contraction type at lower intensities.

## Methods

### Participants

We recruited 33 (17M; 16F) recreationally active, healthy young adult males and females (sex assigned at birth; age 20-28 years). All participants provided written, informed consent and the study was approved by the Hamilton Integrated Research Board (HiREB Project #8244). Participants were non-smokers, and had no previous history of cardiovascular, cerebrovascular, respiratory, autonomic, or neurological disorders. *Table 1* illustrates sex stratified demographic data as well as sex and exercise protocol stratified values for resting physiological measures.

**Table 1.**
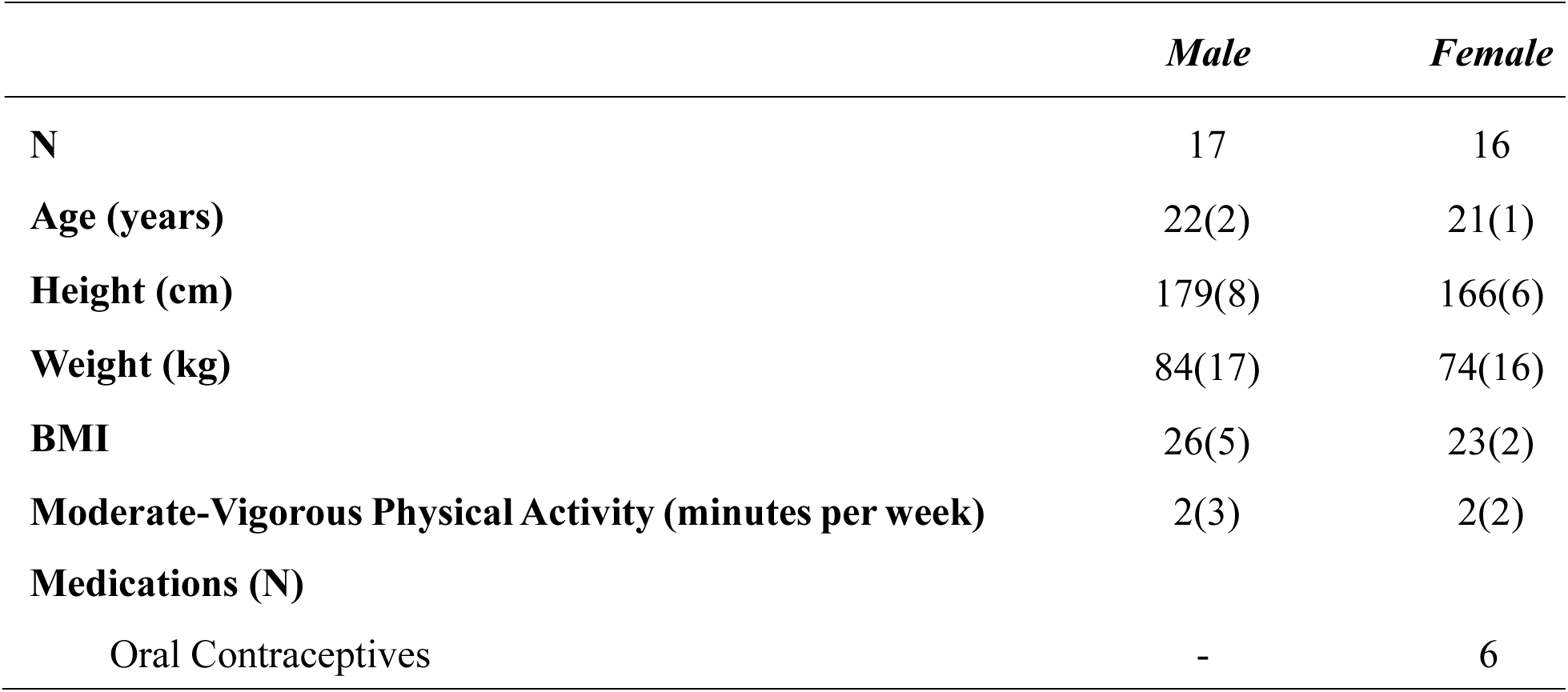
Participant characteristics. Values are reported as means (SD). BMI; body mass index.

**Table 2.** Sex differences for baseline supine systemic and cerebral hemodynamic measurements prior to the start of each HGE protocol. Values are reported as means (SD). Bold values represent statistically significant difference in values between sexes at baseline.

|  | *Male* | *Female* | *p* |
| --- | --- | --- | --- |
| **<u>Heart Rate (BPM)</u>** |  |  |  |
| Static 15% MVC | 63(11) | 67(9) | 0.3 |
| Static 30% MVC | 66(11) | 68(8) | 0.4 |
| Rhythmic 30% MVC | 64(8) | 69(9) | 0.1 |
| Rhythmic 60% MVC | 65(12) | 68(10) | 0.5 |
| **<u>Systolic Blood Pressure (mmHg)</u>** |  |  |  |
| Static 15% MVC | 123(13) | 109(8) | **<0.001** |
| Static 30% MVC | 121(11) | 110(10) | **0.003** |
| Rhythmic 30% MVC | 122(9) | 112(11) | **0.005** |
| Rhythmic 60% MVC | 122(14) | 111(11) | **0.02** |
| **<u>Diastolic Blood Pressure (mmHg)</u>** |  |  |  |
| Static 15% MVC | 68(8) | 72(7) | 0.1 |
| Static 30% MVC | 69(6) | 72(7) | 0.3 |
| Rhythmic 30% MVC | 69(5) | 71(9) | 0.3 |
| Rhythmic 60% MVC | 68(8) | 74(8) | **0.04** |
| **<u>Mean Arterial Pressure (mmHg)</u>** |  |  |  |
| Static 15% MVC | 82(9) | 83(7) | 0.7 |
| Static 30% MVC | 83(7) | 83(6) | 0.8 |
| Rhythmic 30% MVC | 83(3) | 83(8) | 0.9 |
| Rhythmic 60% MVC | 82(9) | 85(8) | 0.3 |
| **<u>Middle Cerebral Artery Blood Velocity (cm/s)</u>** |  |  |  |
| Static 15% MVC | 63(13) | 77(17) | **0.02** |
| Static 30% MVC | 59(13) | 75(18) | **<0.01** |
| Rhythmic 30% MVC | 61(14) | 72(18) | 0.06 |
| Rhythmic 60% MVC | 61(15) | 78(16) | **<0.01** |
| **<u>Middle Cerebral Artery Cerebrovascular Conductance Index (cm/s/mmHg)</u>** |  |  |  |
| Static 15% MVC | 0.8(0.2) | 0.9(0.2) | **0.01** |
| Static 30% MVC | 0.7(0.2) | 0.9(0.2) | **0.01** |
| Rhythmic 30% MVC | 0.7(0.2) | 0.9(0.2) | **0.04** |
| Rhythmic 60% MVC | 0.7(0.2) | 0.9(0.2) | **0.02** |
| **<u>Internal Carotid Artery Diameter (cm)</u>** |  |  |  |
| Static 15% MVC | 0.49(0.05) | 0.46(0.04) | 0.2 |
| Static 30% MVC | 0.49(0.06) | 0.48(0.06) | 0.6 |
| Rhythmic 30% MVC | 0.49(0.06) | 0.48(0.05) | 0.4 |
| Rhythmic 60% MVC | 0.50(0.05) | 0.47(0.04) | 0.2 |
| **<u>Internal Carotid Artery Blood Velocity (cm/s)</u>** |  |  |  |
| Static 15% MVC | 17(5) | 22(5) | **0.02** |
| Static 30% MVC | 16(4) | 22(5) | **<0.01** |
| Rhythmic 30% MVC | 17(5) | 21(3) | **<0.01** |
| Rhythmic 60% MVC | 18(4) | 22(5) | **<0.01** |
| **<u>Internal Carotid Artery Blood Flow (mL/min)</u>** |  |  |  |
| Static 15% MVC | 191(50) | 225(64) | 0.1 |
| Static 30% MVC | 184(48) | 241(85) | **0.03** |
| Rhythmic 30% MVC | 192(71) | 228(51) | 0.1 |
| Rhythmic 60% MVC | 207(47) | 239(61) | 0.1 |
| **<u>Internal Carotid Artery Shear Rate (1/s)</u>** |  |  |  |
| Static 15% MVC | 282(90) | 374(109) | **0.02** |
| Static 30% MVC | 270(86) | 361(98) | **0.01** |
| Rhythmic 30% MVC | 274(100) | 362(73) | **<0.01** |
| Rhythmic 60% MVC | 293(78) | 381(89) | **<0.01** |
| **<u>End tidal CO<sub>2</sub> (mmHg)</u>** |  |  |  |
| Static 15% MVC | 38(3) | 36(3) | 0.3 |
| Static 30% MVC | 38(4) | 36(2) | 0.3 |
| Rhythmic 30% MVC | 37(3) | 36(3) | 0.5 |
| Rhythmic 60% MVC | 37(3) | 36(5) | 0.4 |

### Study Design

The present study consisted of a single visit for both female and male participants. Participants were asked to arrive at least 6 hours fasted as well as having abstained from caffeine for 6 hours, and alcohol and strenuous exercise for 24 hours prior to the protocol. Female participants were tested during the early follicular (or placebo for oral contraceptive users) phase of their menstrual cycle (days 1 to 5 following onset of menses or during placebo pills). The order of each HGE protocol was completed in a randomized order: 1) Static handgrip exercise at 15% maximal voluntary contraction (MVC) (S15), 2) static at 30% MVC (S30), 3) rhythmic handgrip protocol at 30% MVC (R30), and 4) rhythmic at 60% MVC (R60). Each HGE protocol was separated by a 15-minute rest period (20). During each HGE protocol, continuous measures of heart rate (HR), systolic blood pressure (SBP), diastolic blood pressure (DBP), mean arterial pressure (MAP), MCA measures (MCAv, MCA_CVCi_), ICA measures (diameter, mean blood velocity, mean blood flow, and shear rate), P_ET_CO_2_, and Rating of Perceived Exertion (RPE) were collected. Prior to starting the protocol, the participants conducted three maximal handgrip contractions (∼1-2 seconds in duration), separated by 1-minute (20,21), using a handgrip-dynamometer (BMS-G200 Handgrip Dynamometer, Biometrics Ltd, Hamilton, Ontario). The highest force generated of the three maximal voluntary contractions was determined as an individual’s relative MVC. Afterwards, the participant was given a 15-minute rest prior to beginning the experimental protocol.

### Experimental Protocol

Participants were asked to familiarize themselves with the RPE scale, then lie in a supine position during instrumentation. The same sonographer (HAI) insonated and optimized the MCA blood velocity waveform in the contralateral MCA to participant’s dominant (exercise) arm. After the 15-minute rest period, manual brachial artery blood pressure was collected and used to calibrate the continuous blood pressure measurements from the human measured using arterial volume clamping (Human Non-Invasive Blood Pressure Monitor; AD Instruments, Colorado Springs, USA) on the non-dominant hand. Following manual brachial blood pressure measurement, the same experienced sonographer (YM) would secure a stable image and blood velocity signal of the ICA using duplex ultrasonography. The total time for each HGE paradigm was seven minutes with baseline values measured in the first minute, the handgrip exercise in the next five minutes, and recovery data collected in the final minute. RPE was collected 10 seconds before the beginning of each HGE protocol (∼50 sec after the baseline values), at the halfway mark of the HGE protocol, and 10-seconds prior to the cessation of the HGE. For each protocol, a target MVC range was set (+/-5% target MVC) and was displayed to the participant on a monitor to provide visual feedback during exercise. Static HGE was performed as contracting and maintaining the contraction at the target intensity for the entire 5-minute duration of the HGE, whereas rhythmic HGE was performed using a metronome with a 4 second duty cycle (2 second contraction/release interval) at the target intensity.

### Physiological Measurements

Electrocardiogram electrodes were placed in lead II configuration (Bioamp, ML132; ADInstruments) to measure HR. Beat-by-beat arterial pressure (SBP, DBP and MAP) and CO was measured using arterial volume clamping (Human Non-Invasive Blood Pressure Monitor Nano; AD Instruments, Colorado Springs, USA) on the middle finger of the non-dominant hand for all participants. A 2-MHz transcranial Doppler ultrasound (TCD) probe attached to a TCD headset (Multigon Industries, Inc., Elmsford, NY, USA) was placed on the temporal bone contralateral to the exercising limb for all participants to measure MCAv. Once the optimal MCAv waveform was insonated at a depth of 45-60mm, the probe was fixed in place for the duration of the protocol. Mean MCAv was calculated from the peak velocity envelope trace within each protocol. P_ET_CO_2_ was sampled at the mouth via a mouthpiece and gas analyser (ML206; ADInstruments, Colorado Springs, CO, USA). The gas analyser was calibrated to a known gas concentration before each experiment. All physiological data were acquired continuously using an analog-to-digital converter (Powerlab/16SP ML 880; ADInstruments, Colorado Springs, CO, USA) interfaced with a personal computer and sampled at 1KHz. Commercially available software was used to analyse cardiovascular variables (LabChart V7.1; ADInstruments).

Structural images and blood velocity of the ICA were acquired and measured using a 10MHz linear array probe on a high-resolution duplex ultrasound (Terason 3300; Teratech, Burlington, MA, USA). The same experienced ultrasound sonographer (YM) held the position of the ultrasound probe for the duration of each exercise protocol. ICA diameter and blood velocity were recorded continuously. ICA diameter, envelope velocity (i.e., peak velocity), blood flow (half of time-averaged envelope velocity x ICA lumen cross-sectional area), and shear rate analyses were performed using edge-detection software that integrated synchronous diameter and velocity measurements at 190 Hz described in more detail elsewhere (22).

### Data analysis

Cardiovascular, respiratory, and cerebrovascular baseline (BL) and recovery (REC) data were averaged over a 1-minute period immediately before (BL) and after (REC) the HGE, respectively. During HGE, all data were averaged over the final 30 seconds of each minute of the 5-minute protocol at times, 90–120 seconds (time point 1), 150–180 seconds (time point 2), 210–240 seconds (time point 3), 270–300 seconds (time point 4), and 330–360 seconds (time point 5).

### Statistical analysis

All statistical analyses were conducted in the freely available statistical software R (https://cran.r-project.org) using the *lme4* package (https://cran.r-project.org/web/packages/lme4/index.html) to implement the linear mixed effects models.

In all analyses, we tested the interaction between Time (BL, minutes 1-5, REC) and Exercise Protocol (S15, S30, R30, R60) on outcomes of MCAv, MCA_CVCi_, ICA diameter, ICA blood velocity, ICA blood flow, ICA shear rate, HR, SBP, DBP, MAP, CO, and P_ET_CO_2_. Using linear mixed effects models, we accounted for differences at the individual level by fitting individual participants as random effects. Biological sex was included as a fixed effect for all models involving system hemodynamic and respiratory measures (*Equation 1*), and P_ET_CO_2_ was included as a fixed effect in models related to cerebrovascular measures (ie., ICA and MCA measures; *Equation 2*). When omnibus tests detected significant interaction effects between time and exercise protocol, Bonferroni corrected *post-hoc* pairwise comparisons (paired samples t-tests) at each time point were conducted to determine which exercise protocols were significantly different from each other and at which time points. All values are reported as mean (SD) unless otherwise stated. Changes from baseline are represented visually in embedded figures (top right corner) for each variable for interpretation and visualization purposes.

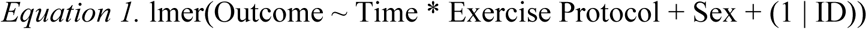

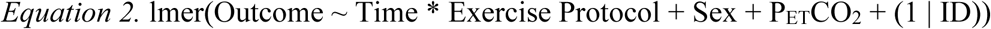

We performed a simulation-based power analysis using the ‘simr’ package in R (https://cran.r-project.org/web/packages/simr/index.html) to detect the minimum physiologically meaningful difference in MCAv (5cm/s) across exercise protocols. The planned linear mixed-effects model included fixed effects of Time, Protocol, their interaction (Time x Protocol), and Sex, and a random intercept for participants as per *Equation 1*. Simulated datasets reflected baseline sex-specific MCAv values of 59.5±10.3cm/s for males and 70.5±12.7 cm/s for females based on references ranges from Alwatban *et al* (23). Between-subject variability was incorporated via random intercepts, and residual variability was sex-specific (SD=10.3cm/s for males, 12.7cm/s for females). Our simulation-based approach determined that a sample size of n=32 was necessary to be 80% powered to detect our *a-priori* effect of interest.

## Results

### Participant characteristics

A total of 33 (17 males; 16 females age 22(1) years; height 173(9)cm; body mass 74(16)kg; body mass index 24(4)kg/m^2^) participants were enrolled into the present study. All participants enrolled in the study identified as cis-gender. No participants were taking medications at time of recruitment aside from oral contraceptives (N=6). One female participant had a copper intrauterine device at the time of enrolment.

### Effects of handgrip protocol on MCA blood velocity and MCA cerebrovascular conductance index

Linear mixed effects models (*Equation 2*) revealed no time by exercise protocol interaction on MCAv (*p*=0.9; *F=0.6*). There was a main effect of time (*p*<0.001; *F*=8.5), where MCAv increased slightly in 3 of 4 protocols from baseline (unchanged in S15). We observed main effects of exercise protocol (*p*<0.001; *F=22.6*), with largest increases in MCAv (from baseline) occurring in R60 (7% increase in MCAv from BL at minute 4) compared to other protocols (*Figure 1A*). There was no time by exercise protocol interaction on MCA_CVCi_ (*p*=0.7; *F=0.8*). We observed main effects of time (*p*<0.001; *F=7.9*), with MCA_CVCi_ decreasing in both static protocols and in R60 over the duration of the test. There was a main effect of exercise protocol (*p*<0.001; *F=6.4*) on MCA_CVCi_, with MCA_CVCi_ decreasing from baseline in both S15 (6% reduction from BL at minute 4), S30 (10% reduction from BL by minute 4) and R60 (10% reduction from BL at minute 5), and MCA_CVCi_ increasing from baseline in R30 (4% increase from BL at minute 1, followed by a return towards baseline at minutes 2-5) (*Figure 1B)*.

**Figure 1.**
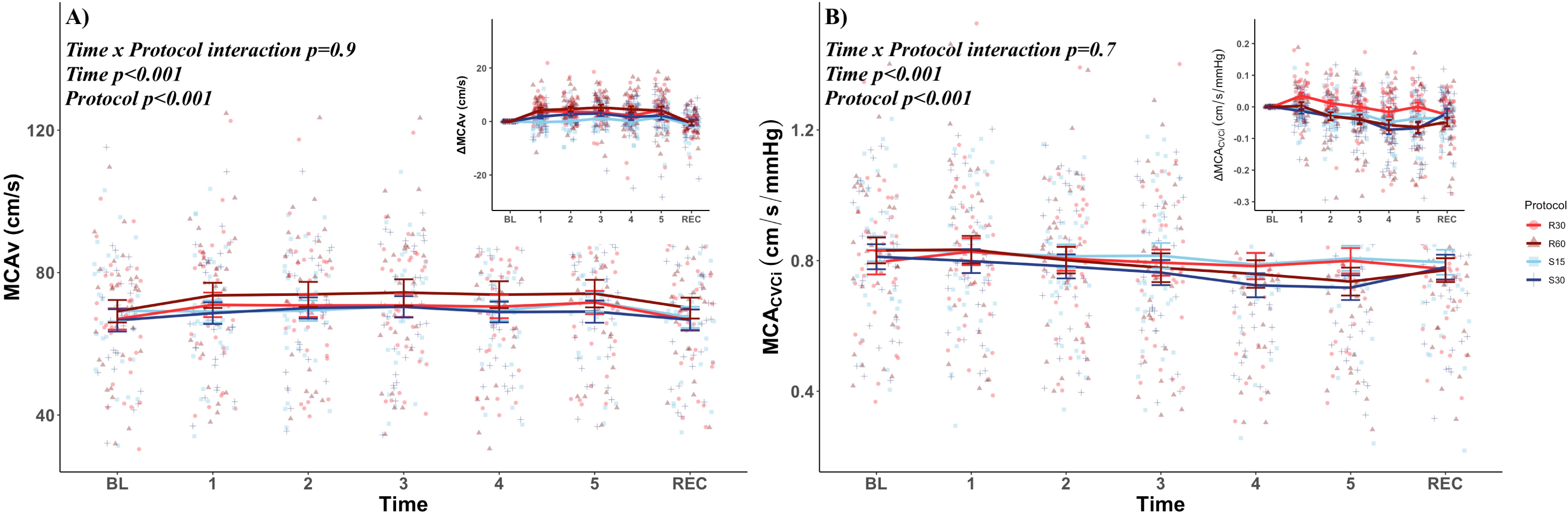
Middle cerebral artery (MCA) responses (MCA blood velocity [MCAv]; MCA cerebrovascular conductance index [MCA_CVCi_]; 1A and 1B, respectively) in N=33 healthy young adults (17 males; 16 females) during different handgrip protocols (R30, R60, S15, S30). Handgrip protocol is plotted by shape and colour (R30 = red circles; R60 = Maroon triangles; S15 = Blue squares; S30 = Navy crosses).

### Effects of handgrip protocol on ICA diameter, blood velocity, blood flow, and shear rate

We observed no time by exercise protocol interaction or main effects of time on ICA diameter (*p*=0.9; *F=0.7; Figure 2A*), blood velocity (*p*=0.5; *F=1.0; Figure 2B*), blood flow (*p*=0.4; *F=1.0; Figure 2C*), or shear rate (*p*=0.7; *F=0.8; Figure 2D*). We identified main effects of exercise protocol for ICA diameter (*p*=0.01; *F=3.9*) and blood flow (*p*=0.001; *F=5.4*), with ICA diameter decreasing from baseline during S30 (4% reduction in diameter from BL at minute 4), and ICA blood flow increased from baseline in R30 (8% increase in ICA blood flow from BL at minute 5). In contrast, R60 was associated with the greatest reduction in ICA blood flow (4% reduction in ICA blood flow from BL at minute 5) relative to other protocols. ICA blood flow responses were similar between both static HGE protocols. There was no effect of exercise protocol on ICA blood velocity (*p*=0.6; *F=0.6*) or shear rate (*p*=0.6; *F=0.6*).

**Figure 2.**
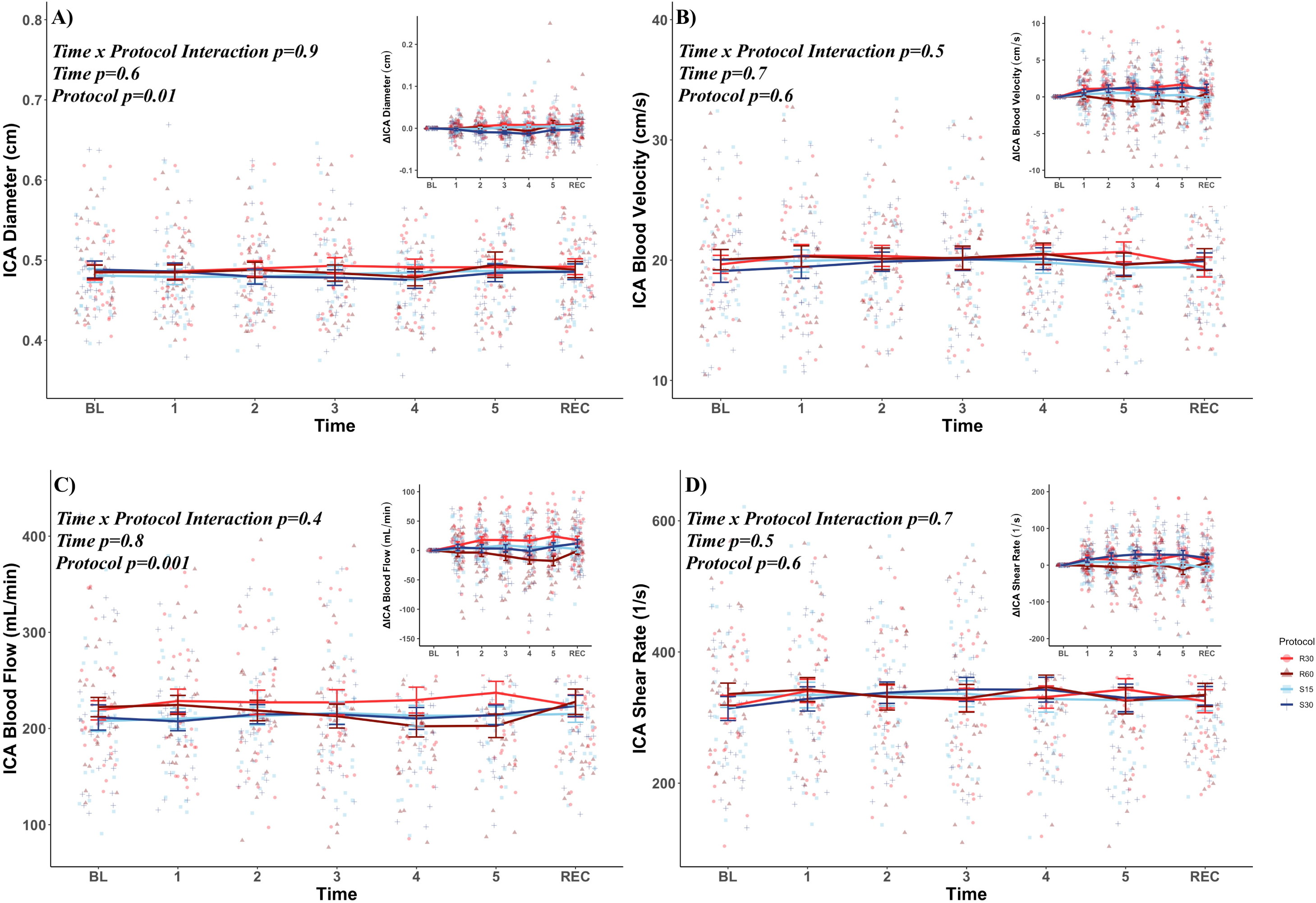
Internal carotid artery (ICA) responses (ICA diameter; ICA blood velocity; ICA blood flow; ICA shear rate; 2A-2D, respectively) in N=33 healthy young adults (17 males; 16 females) during different handgrip protocols (R30, R60, S15, S30). Handgrip protocol is plotted by shape and colour (R30 = red circles; R60 = Maroon triangles; S15 = Blue squares; S30 = Navy crosses).

### Effects of handgrip protocol on systemic physiological measures (heart rate, blood pressure, cardiac output, end-tidal CO_2_)

There was a time by exercise protocol interaction on HR (*p*<0.001; *F=5.4; Figure 3A*), with significantly greater HR responses in higher intensity exercise protocols (S30 [∼17BPM increase from BL to minute 5], R60 [∼13BPM increase from BL to minute 5]) compared to low intensity protocols (S15 [∼5BPM increase from BL to minute 5], R30 [∼4BPM increase from BL to minute 5). There was also a significant time and protocol interaction effect for SBP (*p*<0.001; *F=2.7*), DBP (*p<*0.001; *F=3.4*), and MAP (*p*<0.001; *F=4.5; Figure 3B*). Both S30 (+19mmHg increase from BL to minute 5) and R60 (+15mmHg increase from BL to minute 5) were associated with greater SBP responses compared to S15 (+6mmHg increase from BL to minute 5) and R30 (+6mmHg increase from BL to minute 5). DBP responses followed a similar trend, with increases in DBP occurring in S30 (+13mmHg increase from BL to minute 5) and R60 (+11mmHg increase from BL to minute 5), and little change in S15 (+4mmHg increase from BL to minute 5) and R30 (+5mmHg increase from BL to minute 5).

**Figure 3.**
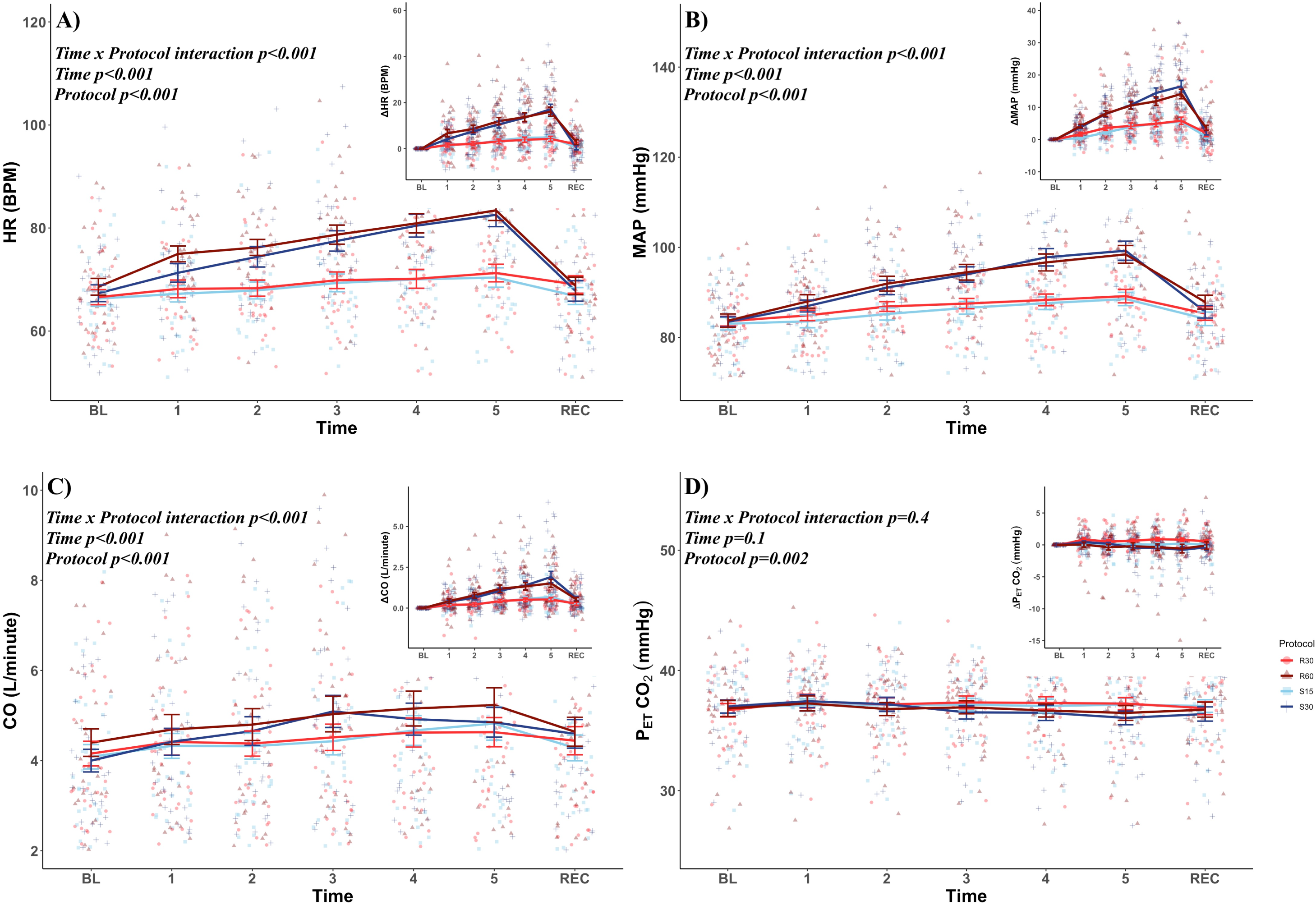
Systemic physiological responses (Heart rate (HR); mean arterial pressure (MAP); cardiac output (CO); end tidal carbon dioxide (P_ET_CO_2_); 3A-3D, respectively) in N=33 healthy young adults (17 males; 16 females) during different handgrip protocols (R30, R60, S15, S30). Handgrip protocol is plotted by shape and colour (R30 = red circles; R60 = Maroon triangles; S15 = Blue squares; S30 = Navy crosses).

We observed significant interaction effects between time and exercise protocol on CO (*p*<0.001; *F=3.7; Figure 3C*). Higher intensity protocols within each contraction type were associated with greater increases in CO (S30: +1.9L/min; R60: +1.5L/min increase from BL to minute 5). The changes in CO in lower intensity protocols were lower, with S15 being associated with a 0.6L/min increase, and R30 being associated with a 0.4L/min increase in CO from BL to minute 5.

There was no time by exercise protocol interaction (*p*=0.4; *F=1.1*) or main effect of time (*p*=0.1; *F=1.7*) on P_ET_CO_2_ (*Figure 3D*). We did observe a main effect of exercise protocol (*p<*0.001; *F=4.7*), with reductions in P_ET_CO_2_ from baseline in both S30 (-1mmHg from BL to minute 5) and R60 (-0.5mmHg from BL to minute 5). There were no changes in P_ET_CO_2_ over the duration of exercise in S15 or R30.

### Rating of perceived exertion during handgrip exercise

There was a time by exercise protocol interaction on RPE (*p*<0.001), such that higher intensities were perceived as more difficult, and this perceived difficulty increased over the duration of the exercise protocol (*Figure 4*). Pairwise comparisons determined that at the halfway point of exercise, higher intensities within each contraction type were perceived as more challenging (S15 vs. S30 *p*<0.001; R30 vs. R60 *p*<0.001). There were no differences between contraction within lower or higher intensities at the halfway mark of exercise.

**Figure 4.**
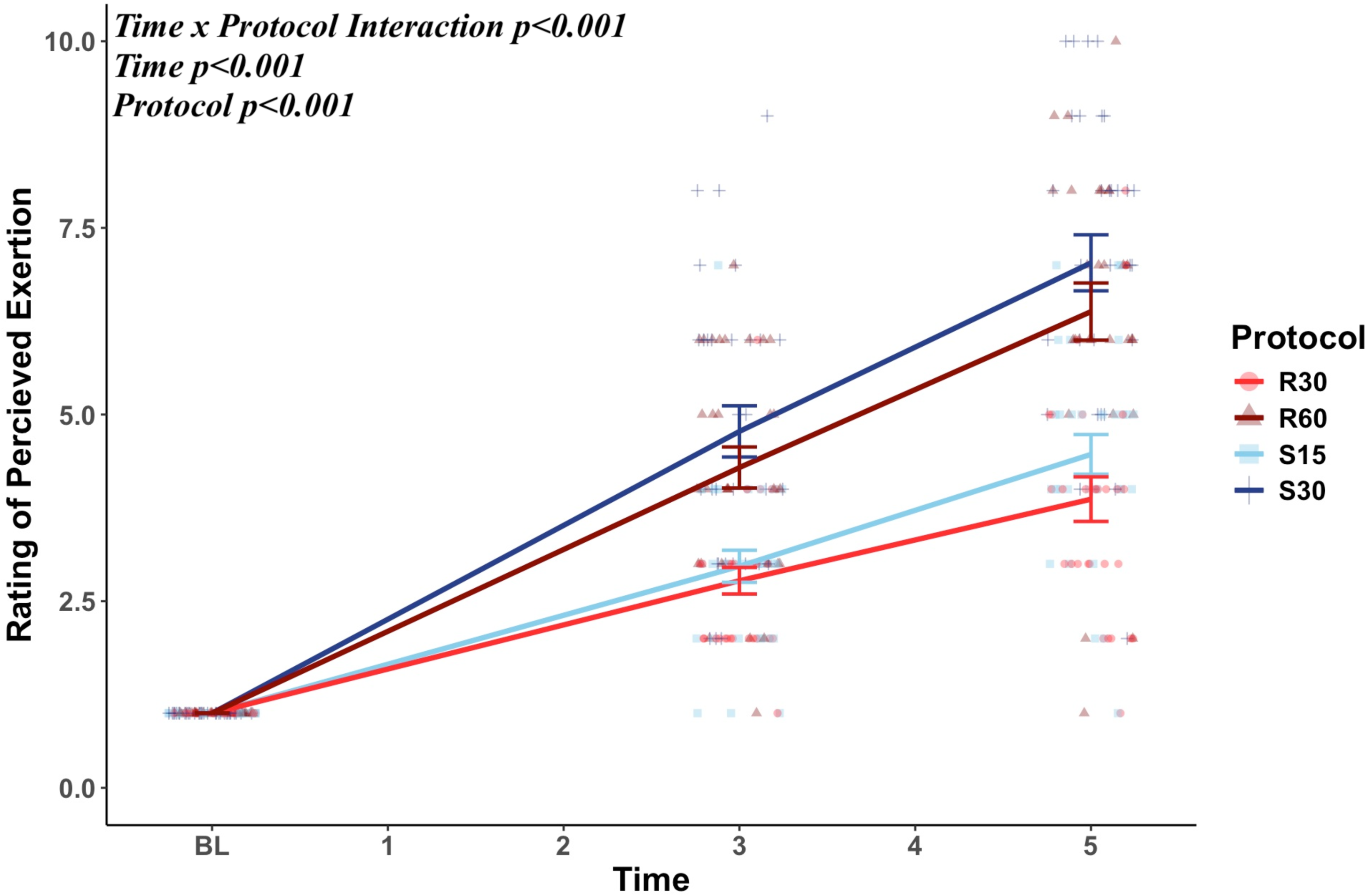
Rating of perceived exertion (RPE) in N=33 healthy young adults (17 males; 16 females) during different handgrip protocols (R30, R60, S15, S30). Handgrip protocol is plotted by shape and colour (R30 = red circles; R60 = Maroon triangles; S15 = Blue squares; S30 = Navy crosses).

## Discussion

To our knowledge, our study is the first to directly compare cerebral and systemic hemodynamic responses across different HGE protocols within the same cohort of participants. In line with our hypothesis, we found greater cerebrovascular responses to rhythmic HGE protocols compared to static, and greater peak systemic cardiovascular responses to static HGE protocols compared to rhythmic. Specifically, we observed that increases in both MCA conductance index (MCA_CVCi_) and ICA blood flow were greatest in rhythmic HGE at 30% of MVC and were accompanied by a modest systemic pressor response (BP, HR, CO) compared to other protocols. Despite eliciting greater systemic responses, higher intensities within each contraction type (S30, R60) were associated with blunted ICA blood flow and MCA_CVCi_ responses compared to lower intensity protocols (S15, R30).

An additional novel component is that we had a sex-balanced sample and conducted exploratory sex-stratified analyses for cerebrovascular (*Supplemental Figures 1 and 2*) and systemic physiological responses (*Supplemental Figures 3*) for future hypothesis generation. Sex differences in neuro-hemodynamic coupling exist, such that there is no correlation between increases in sympathetic outflow and total peripheral resistance in females compared to males (24–26). Visual inspection of our sex-stratified data (provided in *Supplemental Materials*) highlights a greater pressor response in male subjects than in females (*Supplemental Figure 3B, 3F*) though overall patterns of responses between sexes were similar.

### Cerebrovascular Responses to Varying Handgrip Protocols

In our study we observed significant increases in mean MCAv over the duration of both rhythmic HGE protocols, but not in static HGE protocols. This finding aligns with previous work where metrics of CBF did not change across static HGE protocols (27–29). The blunted MCAv response in static HGE supports our hypothesis, and may be linked to greater metaboreflex activation (due to reduced metabolite clearance compared to rhythmic exercise) (17). Static exercise and the accumulation of exercise by-products stimulate afferent nerve fibres and increase MSNA via the muscle metaboreflex (30). Typically, augmented sympathetic outflow during static HGE would be accompanied by an increase in total peripheral resistance and a subsequent rise in blood pressure (26). However, we did not observe pressor differences between static and rhythmic HGE protocols. The observed lack of differences in systemic hemodynamic responses between static and rhythmic HGE protocols is difficult to reconcile based on the available literature, as much of the available literature points to static HGE eliciting greater increases in MSNA compared to rhythmic HGE (11).

Although our interpretation of the role of MSNA in pressor responses is limited given that we did not collect MSNA in our study, we turn to existing literature for context. At the workloads investigated in this study, previous work has shown that static HGE at 30-35% of MVC is associated with a ∼50-100% increase in MSNA and rhythmic HGE at 60% of MVC (11,31–38) is associated with a peak increase of ∼50% in MSNA (39). Given the numerous studies reporting physiological responses to static HGE at 30% MVC (31–38), and the wide range of the sympathetic responses in the literature, it is possible that in our sample individuals experienced less sympathetic outflow during static HGE (lower end of the 50-100% increase in MSNA spectrum) leading to a similar pressor response to R60. In support of this, work from Incognito *et al* that aimed to investigate the inter-individual variability in sympathetic outflow during static HGE at 30% MVC reported significant dispersion across participants, with ∼25% of subjects exhibiting reductions, 34% exhibiting no change, and 41% exhibiting increases in MSNA in the first minute of static HGE, but increased later in exercise (after minute 3 of exercise) in most participants (40). The variable sympathetic response across individuals may be related to differences in the latent period between exercise onset and metabolite production. Indeed, exercise-induced lactate production at a given workload has been shown to be significantly lower in trained individuals compared to their pre-training values (41). Short-term (10 days) exercise training has also been associated with significantly greater capacity for metabolite clearance during exercise (42). Thus, individual levels of fitness certainly have a role in the inter-individual variability in stimulation of the metaboreflex and the subsequent pressor responses. Given that our sample was classified as recreationally active, and we did not collect cardiorespiratory fitness measures, individual differences in fitness and the specifics of exercise habits at the individual level may still partially explain the similar pressor responses between static and rhythmic HGE compared to the available literature. Future studies should aim to investigate whether fitness levels and training modalities direct effects metaboreflex-mediated sympathetic outflow and neurohemodynamic responses.

Verbree *et al* reported vasoconstriction of the MCA via MRI during rhythmic HGE at 60% of MVC (21), which aligns with our finding of a significant reduction in blood flow in the ICA and a drop in MCA conductance during R60. Our findings contradict work from Tarumi *et al,* who observed significant increases in ICA blood flow from baseline in a rhythmic HGE protocol which started at 60% of MVC for the first minute after which intensity was reduced to 40% of MVC for the remainder of the exercise protocol (7 minutes) (43). It is important to note that the work from Tarumi utilized phase-contrast MRI and ICA blood flow was quantified using blood stroke volume instead of blood velocity. There is a possibility for discrepancy between these ultrasound and MRI approaches when quantifying CBF, particularly during dynamic stimuli like exercise (44). Given that we did not observe differences in pressor responses between the two types of HGE, we believe the reduction in ICA blood flow observed in R60 was primarily driven by a drop in ICA blood velocity and increased cerebrovascular resistance (via constriction of MCA; based on Verbree *et al* (21)) whereas the reduction in ICA blood flow in S30 was more likely a product of the observed reduction in ICA diameter rather than constriction of the MCA. The proposed absence of MCA vasoconstriction to static HGE exercise is supported by recent work from Tymko *et al,* as they observed no change from baseline in cerebral noradrenaline spillover (an index of sympathetic outflow) during static HGE at 30% MVC (45), despite consistent reports of static HGE being associated with greater systemic sympathetic outflow (as measured by MSNA).

These findings highlight the complexity of the sympathetic nervous system’s role in cerebrovascular regulation during exercise. Based on the findings from Verbree *et al* and our work, R60 appears to elicit the greatest constrictor (Verbree *et al*) and hemodynamic (our study) response in the MCA but does not affect extracranial arteries like the ICA in a similar manner. An important consideration in the interpretation of the data from Verbree *et al* is the absence of ventilatory measures during exercise, which further limits interpretation of the observed MCA vasoconstriction in their work (21), given the potent role of P_ET_CO_2_ on MCA hemodynamics. Although we found a statistically significant effect of HGE protocol on MCAv, our findings are unlikely to be explained by changes in ventilation, as we observed physiologically negligible (≤1mmHg) differences in P_ET_CO_2_ from baseline in any of the exercise protocols. Given that the observed decreased MCA conductance index and marked reductions in ICA blood velocity and flow in our study support the work from Verbree *et al*, it is possible that their findings of MCA vasoconstriction were similarly not related to exercise-induced hyperventilation.

An alternative explanation for the stability of the MCAv response during S30 compared to R60 despite both demonstrating a reduction in ICA blood flow from baseline and similar MAP responses is the triggering of autoregulatory mechanisms, highlighted by a greater reduction in MCA conductance in S30 compared to R60. Cerebral autoregulation is suggested to be ineffective at frequencies beyond 0.2Hz (5 second cycle) contraction cycles (46). Though speculative, the 4 second duty cycle (2 second contraction, 2 second release) in R60 may have occurred too quickly for autoregulatory mechanisms to elicit the same microvascular myogenic response as S30, despite a similar overall pressor response. Perhaps protection of the cerebral vasculature via buffering of increased systemic driving pressures occurs through distinct mechanisms between static and rhythmic exercise, wherein the former is able to elicit dynamic cerebral autoregulatory mechanisms (ie., microvascular myogenic tone (1)) and the latter acts through *α* –adrenergic receptor mediated vasoconstriction of larger intracranial arteries (47).

The differences in cerebral hemodynamic responses between rhythmic and static HGE are further complicated by typical regulatory variables (MAP, P_ET_CO_2_, CO) being largely similar across conditions at lower and higher workloads, further supporting a difference in local cerebrovascular regulation during rhythmic vs. isometric muscular contractions. The underlying mechanisms for the differential MCAv responses between HGE protocols were not due to systemic cardiorespiratory variables and thus warrants more detailed future investigations using additional physiological parameters such as MSNA, electromyography, and quantification of metabolite concentrations from exercising muscle, or using alternative imaging techniques such as MRI alongside quantification of systemic physiological variables to aid in interpretation.

### Implications for Adapted Exercise Prescription

The primary purpose of this study was to determine cerebral hemodynamic responses to various HGE protocols, with implications for future studies focusing on exercise prescription in populations where dynamic full body exercise may be difficult. In theory, protocols eliciting the greatest increase in cerebral perfusion and vascular shear stress would be linked to the greatest cerebrovascular adaptation if conducted routinely (2). In our work, we found that R30 elicited the greatest increase in MCA_CVCi_ and ICA blood flow, R60 elicited the greatest increase in MCAv, while there were no statistically significant differences in ICA shear rate between protocols. Though increases in perfusion is the primary chronic target of exercise induced vascular adaptation, it is the accompanying elevated shear stress that drives exercise-induced adaptations to the vascular endothelium (48). While not statistically significant, there was a meaningful increase in shear rate compared to baseline in S30 and R30. Indeed, the peak ICA shear rates observed in our study in S30 (346(105)/s) and R30 (343(104)/s) are similar to the ICA shear rates observed during interval training (∼350/s) and aerobic exercise (∼330/s)(49), suggesting that both protocols can adequately stimulate the cerebral vasculature given that both interval training (50) and aerobic exercise (51) have been shown to elicit positive effects on cerebrovascular function and health.

Considering that the increase in shear rate was greatest in S30, an assumption can be made that this protocol (compared to the others) would be most ideal for eliciting cerebrovascular adaptation. While S30 was mostly tolerable for our healthy young sample, it was still the protocol with the highest RPEs, and it remains unclear whether static HGE at 30% MVC would be tolerable in clinical populations with lower exercise capacities. Despite a slightly lower peak shear rate compared to S30, R30 may instead offer a better balance between tolerability and a meaningful cerebrovascular stimulus for exercise-induced adaptations. It is important to acknowledge that typically the shear stimulus on vascular endothelial cells is likely to increase with higher antegrade blood flow (48), even if the metric of shear rate used in this study was not increased substantially. Further, R30 had the lowest relative RPE scores and modest changes in systemic hemodynamics including HR, MAP, and CO. For individuals with chronic health conditions including cardiovascular disease, the high-grade systemic response to higher intensities observed in this study may be contraindicated in early stages of programming to avoid unnecessary risk (52).

Indeed, in populations where exercise capacity is low and the cerebral vasculature may already be in a compromised state (i.e., cardiovascular disease, stroke), prescribing exercise that could stimulate adaptation in the cerebral arteries while mitigating systemic hemodynamic and physiological stress is an important consideration for patient safety. Thus, when considering safety, exercise capacity, and risk management for acute cardio- and cerebrovascular events, low-moderate intensity rhythmic HGE may be the most efficient point of entry to facilitate improvements in cerebrovascular function and brain health through exercise while limiting the exercise pressor response. While exercise intensity plays a significant role in systemic and cerebrovascular adaptation as well as overall brain health (51), introducing exercise at lower intensities is often the recommended approach with eventual progression in both volume and intensity once exercise capacity has increased (53). Although our results are in support of R30 as an ideal starting stimulus, further interventional work in vulnerable populations is necessary to determine if routine rhythmic HGE exercise done at 30% of MVC can elicit measurable improvements in cerebrovascular (cerebrovascular reactivity, autoregulation, neurovascular coupling) and general neurological health and function (brain structure, MRI, cognitive performance).

### Limitations and Methodological Considerations

While acute responses offer valuable insight into regulation of cerebral hemodynamics during exercise, it remains unclear how these acute responses may translate into long-term adaptations. Interventional work is necessary to determine whether increases in indices related to cerebral blood flow and cerebrovascular conductance are predictive of long-term improvements in cerebrovascular function or cognition. Another limitation for this work is the use of TCD to assess intracranial cerebral hemodynamics, as a constant cross-sectional area is assumed (54), despite evidence that MCA diameter changes with rhythmic HGE (21).

Although TCD-derived indices are limited, our work is strengthened by the inclusion of ICA responses to exercise, providing information related to a principle extracranial conduit artery and aiding in interpretation of MCA data. Another limitation related to cerebrovascular measures in this study is the inclusion of unilateral data for both ICA and MCA, as we could not assess hemispheric differences during exercise. Use of bilateral measurements provide a more complete representation of global cerebrovascular responses.

While no formal sex differences analysis was conducted, we accounted for biological sex in our statistical models and included sex-stratified analyses and figures in the *Supplemental Materials* to support future hypothesis generation. Additionally, we were unable to account for the potential influence of physical fitness or habitual resistance training exposure on cerebrovascular responses. Participants with prior resistance training experience and higher grip strength (55) may have been better able to tolerate HGE, particularly at higher intensities, potentially contributing to inter-individual variability in responses. As such, the heterogeneity in MVC capacity across participants introduces variability that may affect the interpretation of cerebrovascular outcomes. Lastly, future work should explore alternative metrics related to participant perception of tolerability and fatigue during HGE, as in this study we relied solely on self-reported RPE to determine tolerability, which is subjective and thus has inherent limitations related to validity (56). Using RPE alongside more physiological markers including but not limited to microneurography and electromyography could improve understanding of the best balance between tolerability and physiological stimulus with implications on optimizing HGE prescriptions for populations requiring adapted physical activity.

## Conclusion

In summary, our findings suggest that cerebrovascular responses to HGE are variable depending on the protocol administered. Low-moderate intensity rhythmic HGE (ie., R30) elicited the most robust increases in index measures of cerebral blood flow (ICA blood flow) and cerebrovascular conductance (MCA_CVCi_) with minimal systemic cardiorespiratory strain and lower RPEs, suggesting a favorable balance between physiological responses and tolerability. While the underlying mechanisms for differential MCAv responses remain unclear, our data highlight the need for further mechanistic and interventional studies to explore cerebrovascular regulation during HGE and the potential accompanying training adaptations. These results suggest that low-moderate intensity rhythmic HGE may be a promising entry point for exercise prescriptions when the goal is to support cerebrovascular health and function, particularly in populations requiring adapted forms of physical activity.

---

*Supplemental Figure 1.* Sex stratified middle cerebral artery (MCA) responses (MCA blood velocity [MCAv]; MCA cerebrovascular conductance index [MCA_CVCi_]) data in N=33 healthy young adults (17 males [A,B]; 16 females [C,D]) during different handgrip protocols (R30, R60, S15, S30). Handgrip protocol is plotted by shape and colour, with different colour palettes between male and female data.

*Supplemental Figure 2.* Sex stratified internal carotid artery (ICA) responses (ICA diameter; ICA blood velocity; ICA blood flow; ICA shear rate) data in N=33 healthy young adults (17 males [A-D]; 16 females [E-H]) during different handgrip protocols (R30, R60, S15, S30). Handgrip protocol is plotted by shape and colour, with different colour palettes between male and female data.

*Supplemental Figure 3.* Sex stratified (Heart rate [HR]; mean arterial pressure [MAP]; cardiac output [CO]; end tidal carbon dioxide [P_ET_CO_2_]) data in N=33 healthy young adults (17 males [A-D]; 16 females [E-H]) during different handgrip protocols (R30, R60, S15, S30). Handgrip protocol is plotted by shape and colour, with different colour palettes between male and female data.

