## Supplementary figures and images for "Cerebrovascular Responses to Static and Rhythmic Handgrip Exercises"

### Supplemental Figure 1

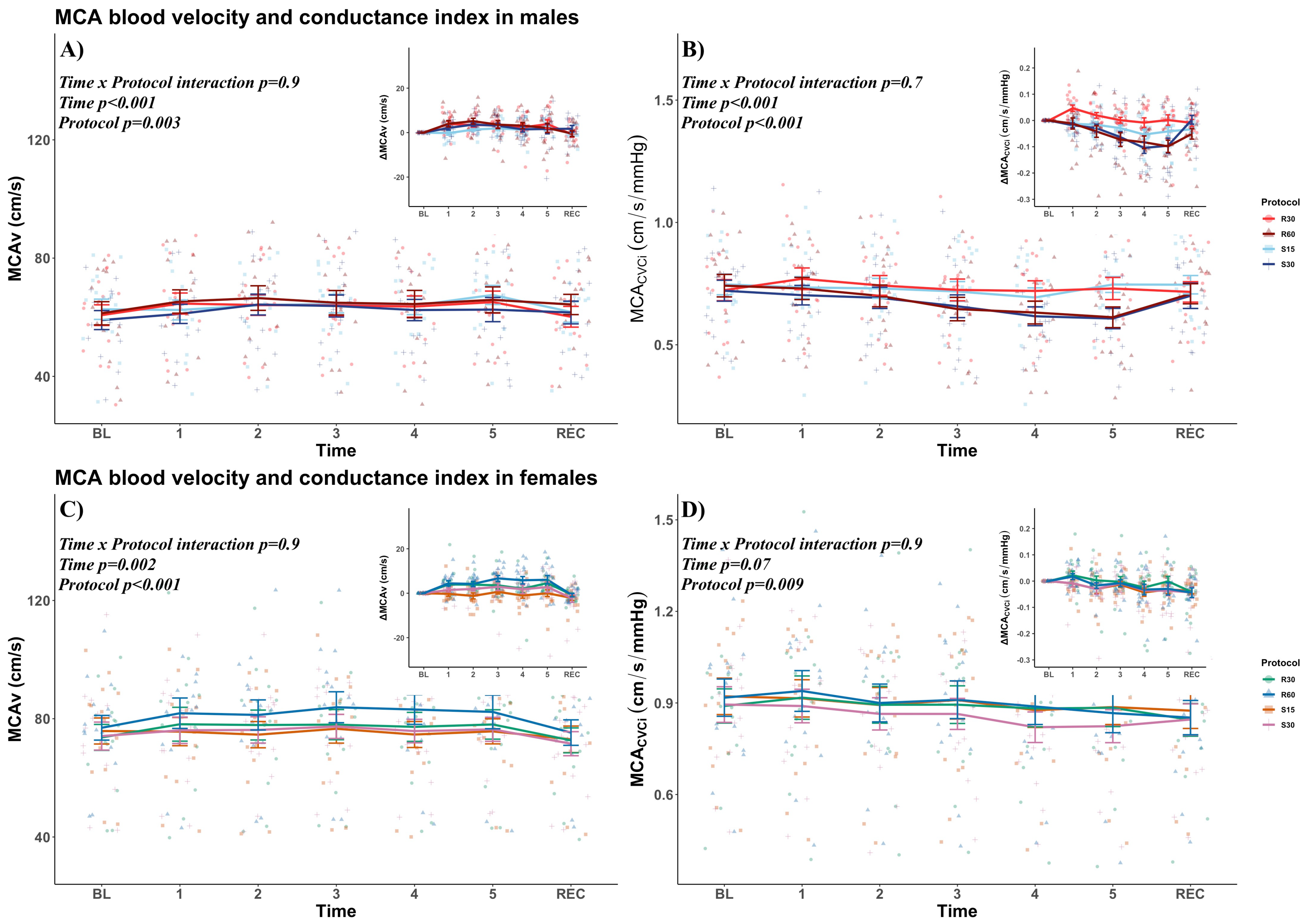

### Supplemental Figure 2

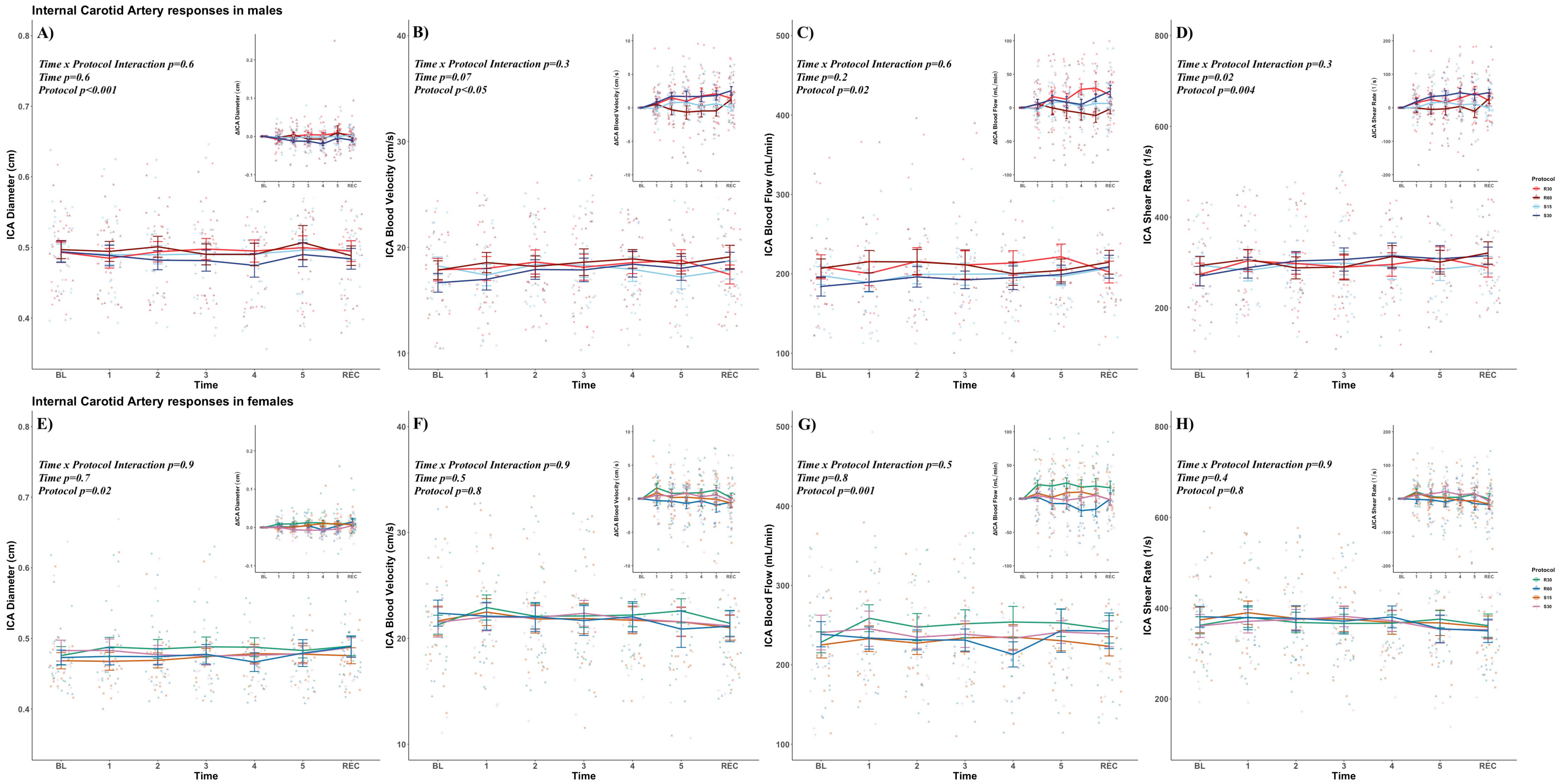

### Supplemental Figure 3

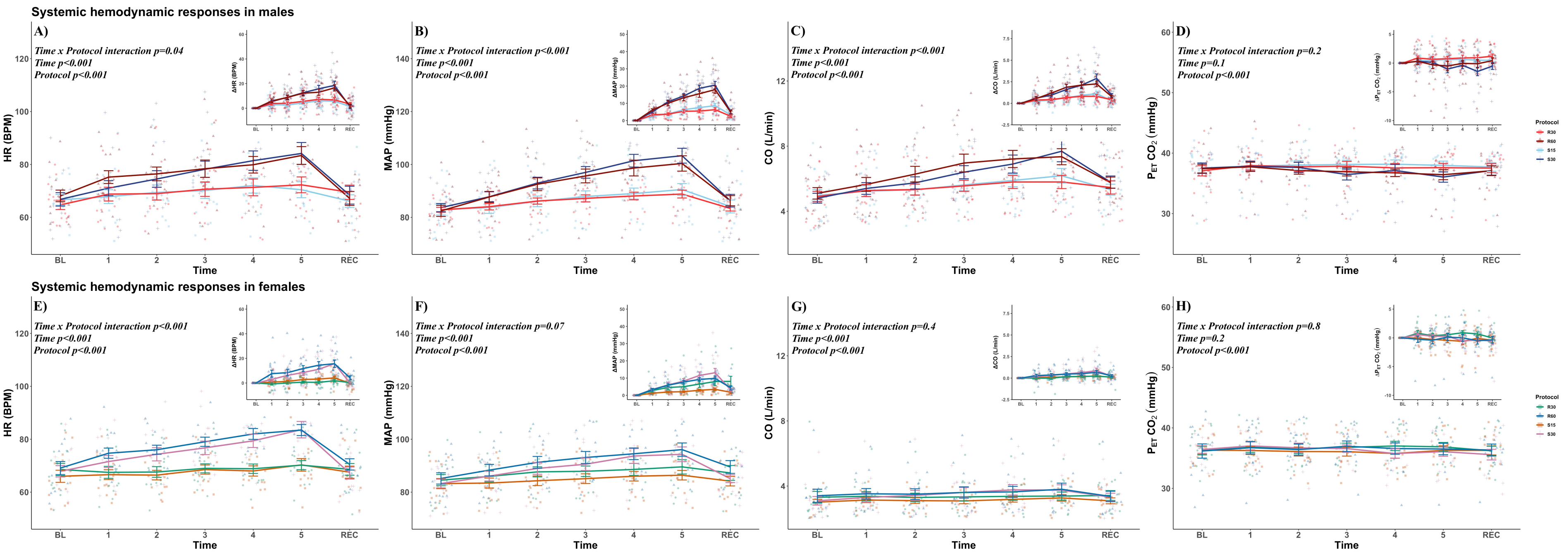
